# A truncated form of the p27 CDK inhibitor translated from pre-mRNA causes G_2_-phase arrest

**DOI:** 10.1101/2022.01.12.476115

**Authors:** Daisuke Kaida, Takayuki Satoh, Ken Ishida, Rei Yoshimoto, Kanae Komori

**Affiliations:** Graduate School of Medicine and Pharmaceutical Sciences, University of Toyama, Toyama 930-0194, Japan; Department of Applied Biological Sciences, Faculty of Agriculture, Setsunan University, Hirakata, Osaka 573-0101, Japan

## Abstract

Pre-mRNA splicing is an indispensable mechanism for eukaryotic gene expression. Splicing inhibition causes cell cycle arrest at G1 and G_2_/M phases, which is thought to be one of the reasons for the potent antitumor activity of splicing inhibitors. However, the molecular mechanisms underlying the cell cycle arrest have many unknown aspects. In particular, the mechanism of G_2_/M-phase arrest caused by splicing inhibition is completely unknown. Here, we found that lower and higher concentrations of pladienolide B caused M-phase and G2-phase arrest, respectively. We analyzed protein levels of cell cycle regulators and found that a truncated form of the p27 CDK inhibitor, named p27*, accumulates in G_2_-arrested cells. Overexpression of p27* caused partial G_2_-phase arrest. Conversely, knockdown of p27* accelerated exit from G2/M phase after washout of splicing inhibitor. These results suggest that p27* contributes to G2/M-phase arrest caused by splicing inhibition. We also found that p27* bound to and inhibited M-phase cyclins, although it is well known that p27 regulates G_1_/S transition. Intriguingly, p27*, but not full-length p27, was resistant to proteasomal degradation and remained in G_2_/M phase. These results suggest that p27*, which is a very stable truncated protein in G_2_/M phase, contributes to G_2_-phase arrest caused by splicing inhibition.

## Introduction

More than 95% of protein-coding genes in humans consist of exons and intervening sequences, namely, introns (1-3). Introns are removed and exons are joined by pre-mRNA splicing to produce mature mRNA: the template for translation. Pre-mRNA splicing is carried out by a macromolecular ribonucleoprotein complex, the spliceosome, which consists of U1, U2, U4/U6, and U5 small nuclear ribonucleoproteins (snRNPs). Therefore, defects in the spliceosome and other splicing factors might cause the accumulation of unspliced or partially spliced mRNAs. If such mRNAs are exported to the cytoplasm and translated into proteins, these proteins would have different amino acid sequences and might have different functions from the original proteins. Such proteins translated from pre-mRNA are supposed to be non-functional or deleterious by inhibiting cellular functions (4) and might cause splicing-related diseases (5-9). To prevent translation from unspliced or partially spliced mRNAs, cells have multiple mRNA quality control mechanisms. For example, the nuclear exosome degrades such unspliced mRNAs (10). Even if unspliced mRNAs escape from this degradation, the export of these mRNAs is strictly prohibited (11-13). If unspliced and partially spliced mRNAs nonetheless leak from the nucleus, they are degraded by nonsense-mediated mRNA decay (NMD) in the cytoplasm (14). These mRNA quality control mechanisms prevent the production of non-functional proteins translated from pre-mRNAs and protect the integrity of the proteome.

To date, several potent splicing inhibitors, including spliceostatin A (SSA) and pladienolide B (Pla-B), have been reported (15, 16). SSA and Pla-B have a molecular target in common, the SF3B complex (a subcomplex of U2 snRNP), their binding to which inhibits pre-mRNA splicing (15, 16). SSA and Pla-B possess very potent anti-tumor activity and cause G_1_- and G_2_/M-phase arrest (15-18). Such cell cycle arrest is thought to be one of the reasons for the antitumor activity of splicing inhibitors. Both compounds weaken the binding between the SF3B complex and pre-mRNA, resulting in splicing perturbation (19, 20). As described above, these compounds share many common features, but there is a distinct difference: SSA binds to its target covalently, but Pla-B binds to it non-covalently (21).

We previously investigated the molecular mechanism of the G_1_-phase arrest induced by splicing inhibition and found that upregulation of the cyclin-dependent kinase (CDK) inhibitor p27 and a C-terminus-truncated form of p27, named p27*, is one of the causes of G_1_-phase arrest induced by splicing inhibition (22, 23). Interestingly, p27* is translated from pre-mRNA of *CDKN1B* despite mRNA quality control mechanisms (15). In addition, we found that the downregulation of cyclin E1, cyclin E2, and E2F1 is another cause of G_1_-phase arrest induced by splicing inhibition (24). As described above, we revealed that splicing inhibition causes G_1_-phase arrest through various mechanisms. However, how splicing inhibitors cause G_2_/M-phase arrest remains completely unclear.

Cell cycle progression from S phase to M phase is tightly regulated by the kinase activity of CDK–cyclin complexes (25, 26). Among CDKs and cyclins, Cdk1 (also known as Cdc2), cyclin A, and cyclin B are critical regulators of G_2_/M phase (27-29). The kinase activity of CDK–cyclin complexes is regulated by several mechanisms. The protein levels of cyclins A and B oscillate during the cell cycle. Cyclin A starts accumulating in S phase and suddenly decreases at early M phase (30-34). Cyclin B starts accumulating in G_2_ phase and decreases at late M phase (28, 35-37). Because cyclin A and B proteins are required for Cdk1 activity, such activity is proportional to the amount of cyclin proteins. In addition to the amount of cyclin proteins, Cdk1 activity is regulated by the phosphorylation status of Cdk1. Cdk1 is phosphorylated at Thr14 and Tyr15 residues in G_2_ phase, which negatively regulates Cdk1 kinase activity (28, 38, 39). At the end of G_2_ phase, Cdk1 is dephosphorylated by Cdc25C for activation, and consequently cells enter M phase (28, 40). In addition to the above two mechanisms, CDK inhibitor proteins negatively regulate the kinase activity of CDK–cyclin complexes (26, 41). As mentioned above, p27 is such a CDK inhibitor that controls G_1_/S transition (42). In fact, p27 is highly expressed in G_0_/G_1_ phase and degraded by the ubiquitin–proteasome pathway from S to M phase (43, 44). Therefore, p27 does not appear to control G_2_/M transition under physiological conditions. However, interestingly, knockout of Skp2, which is a component of SCF (Skp, Cullin, F-box containing complex) ubiquitin ligase for p27 ubiquitylation, causes the accumulation of p27 in G_2_ phase. The accumulated p27 inhibits M-phase cyclins and consequently causes G_2_ arrest (44).

The molecular mechanisms of the G_2_/M-phase arrest caused by splicing inhibition remain unknown. Meanwhile, it is speculated that p27* can escape from proteasomal degradation (15), but there is no experimental evidence in support of this. In this study, we investigated the molecular mechanisms of G_2_/M-phase arrest. Surprisingly, p27*, which is one of the factors behind G_1_-phase arrest, was also shown to contribute to the G_2_/M-phase arrest caused by splicing inhibition. We also found that p27* inhibits M-phase cyclin and is resistant to proteasomal degradation. This study should contribute to a comprehensive understanding of the molecular mechanisms of cell cycle arrest and anti-tumor activity caused by splicing inhibition.

## Results

### Low and high concentrations of Pla-B cause M-phase and G_2_-phase arrest, respectively

To understand the molecular mechanisms of cell cycle arrest upon splicing inhibition, we analyzed the effect of Pla-B on cell cycle progression after cell cycle synchronization by a double thymidine block. We found that 3 or 10 ng/ml Pla-B treatment caused G_2_/M-phase arrest by measuring the DNA content of each cell (Fig. 1A and 1B). Pla-B-treated cells showed an almost identical pattern to MeOH-treated cells until 8 h, suggesting that Pla-B treatment did not affect S-phase progression under these experimental conditions, which is consistent with a previous report (17) (Fig. 1B). We also investigated the morphology of Pla-B-treated cells because the DNA contents of G_2_- and M-phase cells are the same, preventing us from distinguishing G_2_- and M-phase cells based on their DNA content. To this end, we observed the cell morphology and counted the number of cells that were round, a feature of M-phase cells (45). The proportion of round cells among MeOH-treated cells was approximately 40% at 10 h and then decreased (Fig. 1C). Upon treatment with 3 ng/ml Pla-B, round cells had accumulated at 20–36 h (Fig. 1C). In contrast, cells treated with 10 ng/ml Pla-B were flat until 36 h (Fig. 1C). These results suggested that low concentrations of Pla-B caused M-phase arrest, while high concentrations of it caused G_2_-phase arrest.

**Figure 1.**
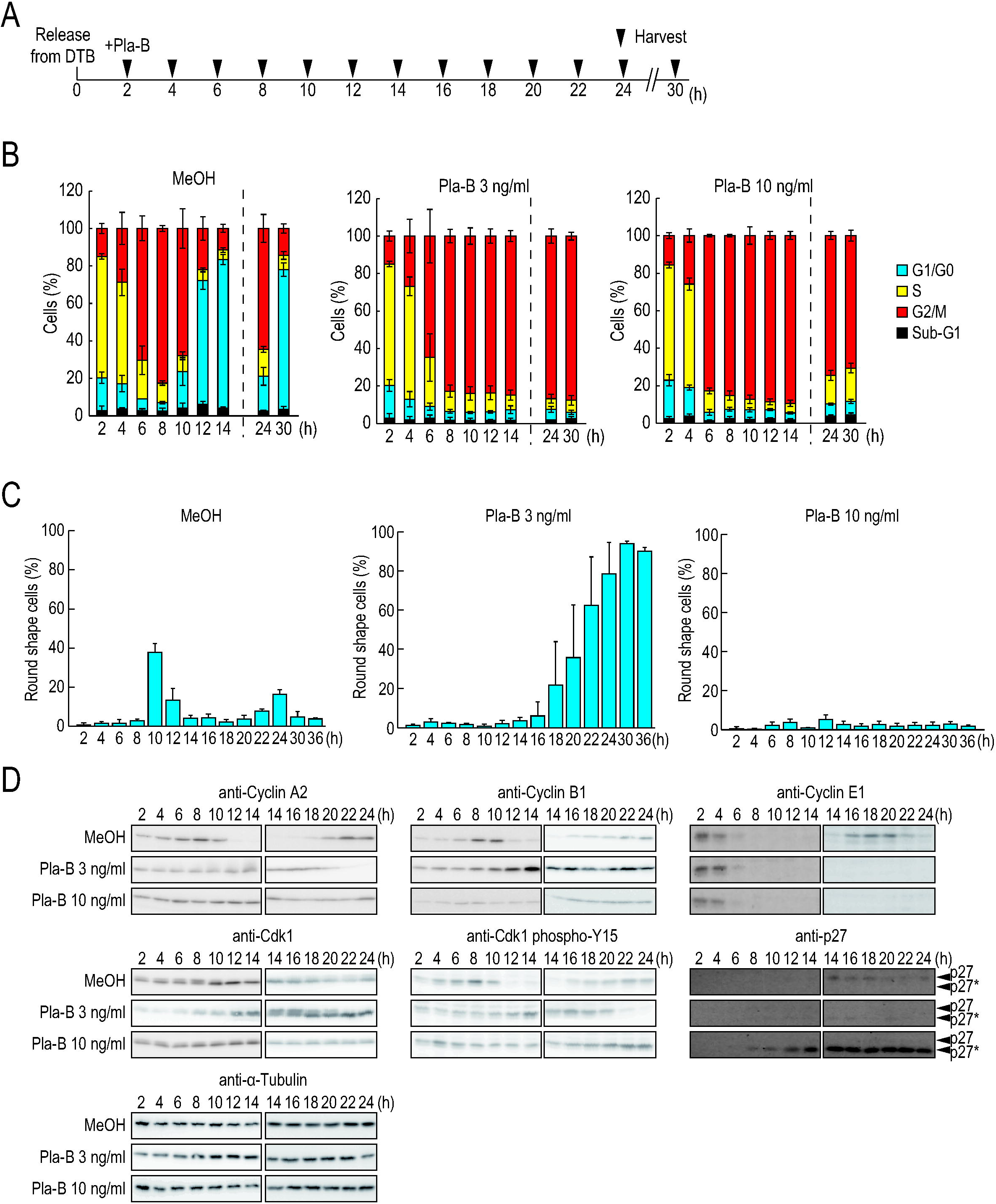
Low doses of Pla-B cause M-phase arrest and high doses of it cause G_2_-phase arrest. (A–D) Two hours after release from a double thymidine block (DTB), synchronized HeLa S3 cells were treated with MeOH or the indicated concentrations of Pla-B. Black triangles indicate the time points of cell harvest and sample preparation (A). Cell cycle was analyzed at the indicated time points by cytometry (B). Morphology of the cells was observed under a microscope and round cells were counted at the indicated time points (C). Protein samples were prepared at the indicated time points. The protein levels of indicated proteins and phosphorylation status of Cdk1 were analyzed by immunoblotting. Protein levels of α-tubulin were analyzed as an internal control (D). Error bars indicate s.d. (n = 3).

In addition to the DNA content and cell morphology, we analyzed the protein levels of cell cycle regulators by immunoblotting upon Pla-B treatment (Fig. 1D). Degradation of cyclin A2, which is a hallmark of M-phase entry, was delayed in cells treated with 3 ng/ml Pla-B (30, 32), and no degradation of cyclin A2 was observed in cells treated with 10 ng/ml Pla-B (Fig. 1D), suggesting that a low concentration of Pla-B delays G_2_/M transition, while a higher concentration of it inhibits this transition. Cyclin B1, which is degraded at late M phase (anaphase) (28, 35-37), was stable in cells treated with 3 ng/ml Pla-B (Fig. 1D), suggesting that a lower concentration of Pla-B causes cell cycle arrest before anaphase. In cells treated with 10 ng/ml Pla-B, cyclin B1 protein levels were lower than those in MeOH-treated cells. This decrease might cause G_2_-phase arrest (see below). We also examined the phosphorylation status of Cdk1 Y15, which is dephosphorylated at the end of G_2_ phase and its dephosphorylation is required for the activation of Cdk1 and G_2_/M transition (40, 46). Dephosphorylation of Cdk1 Y15 was delayed in cells treated with 3 ng/ml Pla-B, and no dephosphorylation of Cdk1 Y15 was observed in cells treated with 10 ng/ml Pla-B (Fig. 1D). These results supported the idea that low concentrations of Pla-B cause M-phase arrest, while high concentrations of it cause G_2_-phase arrest, which is consistent with the above results (Fig. 1B-D). We also analyzed other cell cycle regulators which regulate G1/S transition including p27 and cyclin E1. Surprisingly, we found that a faster migrating isoform of p27, but not full-length p27 (hereafter called p27 FL), accumulated in the cells treated with 10 ng/ml Pla-B that were arrested in G_2_ phase (Fig. 1D). Previously, we reported that a C-terminal-truncated form of p27 translated from *CDKN1B* pre-mRNA, designated p27*, is produced in splicing-deficient cells (15). It was also reported that another shorter form of p27 is produced by caspase cleavage (47). To investigate how the faster migrating isoform of p27 is produced in Pla-B-treated cells, we tested whether poly(ADP-ribose) polymerase (PARP) is cleaved in Pla-B-treated cells because PARP is a well-known caspase substrate (48). We observed the cleaved PARP at 48 h but not at 24 h after Pla-B addition, suggesting that caspase is activated at 48 h but not at 24 h (Fig. 2A). Meanwhile, the faster migrating form of p27 was observed at much earlier time points (Fig. 1D and 2A). Therefore, this faster migrating form might not be the p27 cleaved by caspase. We also used a reporter plasmid to detect the translation of *CDKN1B* pre-mRNA, and found that the pre-mRNA was translated into p27* in Pla-B-treated cells (Fig. 2B and 2C), consistent with our previous results (15). These results suggest that the faster migrating form of p27 is p27* that is translated from pre-mRNA.

**Figure 2.**
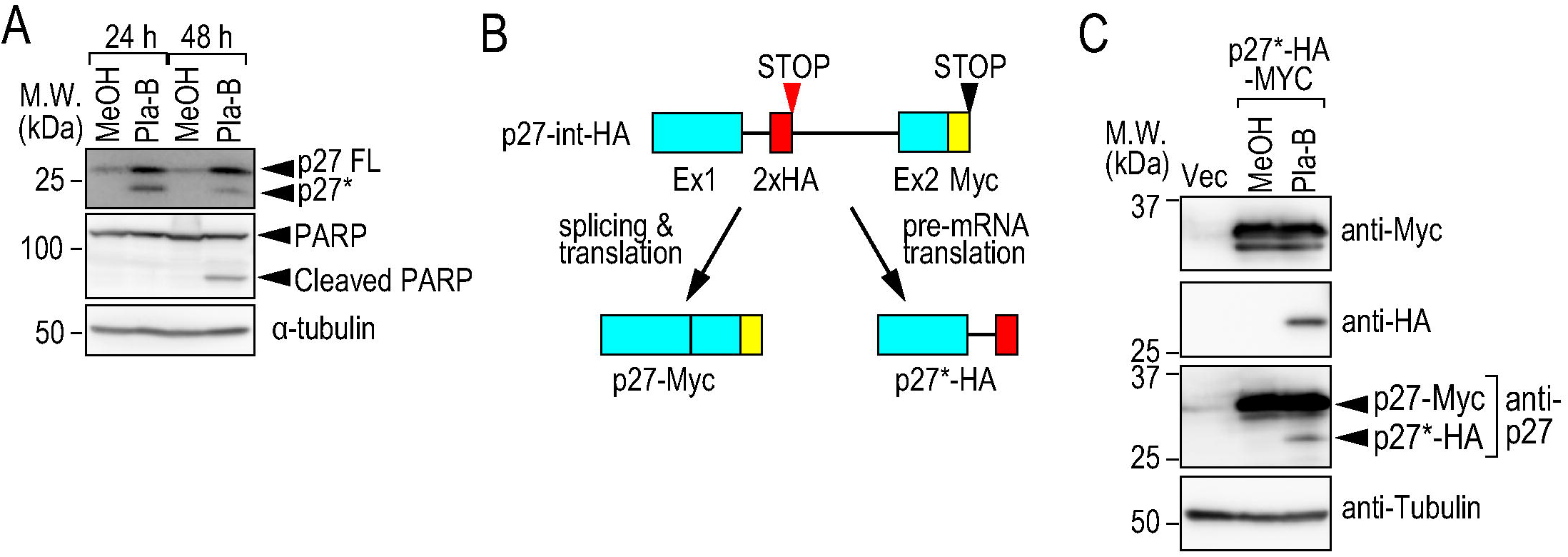
p27* is translated from pre-mRNA. (A) HeLa S3 cells were treated with 10 ng/ml Pla-B for 24 or 48 h, and the amount of indicated protein was analyzed by immunoblotting. (B) The structure of the p27-int-HA reporter gene (15). When the intron sequence is spliced out, the Myc-tagged protein is produced. When splicing is inhibited, the intron sequence is translated to the in-frame stop codon within the intron and the HA-tagged protein is produced. Cyan rectangles, red rectangles, yellow rectangles, and horizontal lines indicate exon sequences, HA tag, Myc tag, and intron sequences, respectively. The red and black triangles indicate the in-frame stop codon within the intron and the bona fide stop codon, respectively. (C) HeLa S3 cells were transfected with vector or p27-int-HA reporter plasmid and treated with 10 ng/ml Pla-B for 24 h. The amount of indicated protein was analyzed by immunoblotting.

### Overexpression of p27* causes G_2_-phase arrest

Because p27* accumulated in the G_2_-arrested cells, we tested whether p27* induces G_2_-phase arrest. At first, we assessed the effect of overexpression of p27 FL and p27* on cell proliferation and cell viability, and found that both forms inhibited cell proliferation at 48 and 72 h after plasmid transfection; however, they did not affect cell viability, suggesting that p27 FL and p27* overexpression causes cell cycle arrest (Fig. 3A–C). Next, we investigated whether overexpression of these forms causes G_2_/M-phase arrest. We transfected non-synchronized cells with p27 FL or p27* expression vector and analyzed the cell cycle. Both p27 FL and p27* caused cell cycle arrest at G_1_ and G_2_/M phases and the proportion of G_2_/M-phase-arrested cells was significantly higher than in thymidine-treated cells, which show G_1_-phase arrest (p-values from one-way ANOVA with Tukey’s test, p27 FL vs. DTB: *p* = 0.00499; p27* vs. DTB: *p* = 0.00314) (Fig. 3D). These results suggest that the overexpression of p27 FL or p27* causes not only G_1_-phase arrest but also G_2_/M-phase arrest, although G_1_-phase arrest is more prominent.

**Figure 3.**
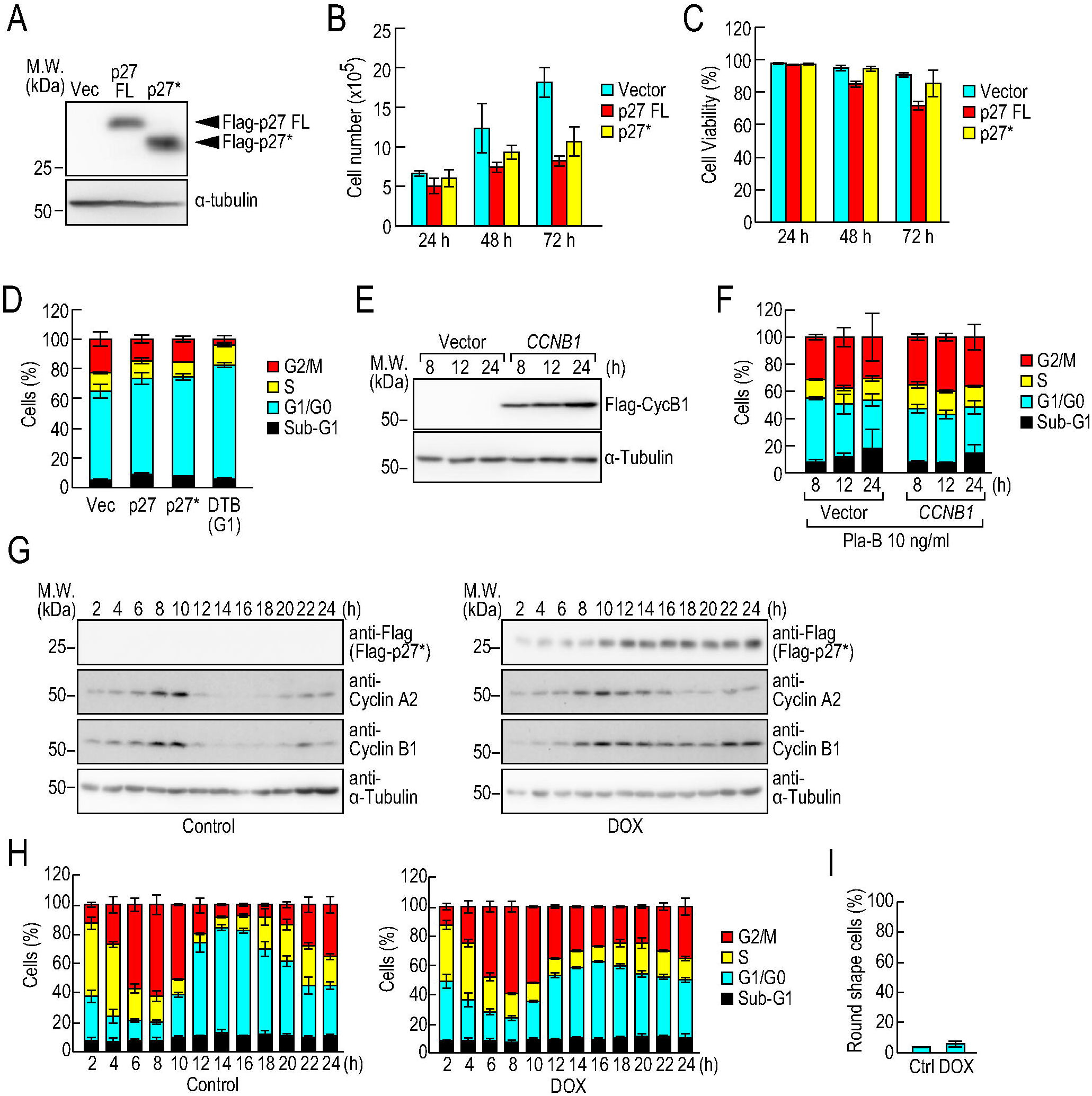
Overexpression of p27* causes partial G_2_-phase arrest. (A–C) HeLa S3 cells were transfected with vector, Flag-p27 FL, or Flag-p27* plasmid. The amount of indicated protein was analyzed by immunoblotting (A). The number of cells (B) and cell viability (C) were analyzed at the indicated time points. (D) HeLa S3 cells were transfected with vector, Flag-p27 FL, or Flag-p27* plasmid. As a control, HeLa S3 cells were synchronized by a double thymidine block. Cell cycle was analyzed by cytometry. (E, F) HeLa S3 cells were synchronized by a double thymidine block and transfected with vector or Flag-*CCNB1* plasmid. The cells were treated with 10 ng/ml Pla-B. The protein levels of the indicated proteins were analyzed by immunoblotting (E). Cell cycle was analyzed by cytometry (F). (G–I) Cells expressing Flag-p27* under the control of tetracycline were synchronized by a double thymidine block. The cells were treated with 2 μg/ml doxycycline (DOX) or DMSO (Control) at the same time as the second thymidine treatment. After release from the double thymidine block, the cells were treated with 2 μg/ml DOX or DMSO and the protein levels of the indicated proteins were analyzed by immunoblotting (G). Cell cycle was analyzed by cytometry (H). The morphology of the cells was observed, and the number of round cells was counted at 20 h after release from the double thymidine block (I).

As described above, the level of cyclin B1 protein decreased in 10 ng/ml Pla-B-treated cells. It is possible that the decrease of cyclin B1 causes G_2_/M-phase arrest. To test this hypothesis, we investigated the cell cycle progression of *CCNB1*-overexpressing cells after Pla-B treatment. The overexpression of *CCNB1* was confirmed by immunoblotting (Fig. 3E). At 8 h after release from double thymidine block, ∼35% of the vector-transfected and Pla-B-treated cells were in G_2_/M phase, and the percentages of the cells were not changed at 12 and 24 h, suggesting that these cells were arrested at G_2_/M phase (Fig. 3F). Notably, we found that a substantial proportion of the cells did not exit G_1_ phase after the double thymidine block and plasmid transfection, presumably because of cell cycle synchronization and transfection stress (22). Indeed, ∼45% of the cells were in G_1_ phase at 8, 12, and 24 h regardless of splicing inhibition (Fig. 3F). We assessed the effect of *CCNB1* overexpression on cell cycle progression of Pla-B-treated cells and found that *CCNB1* overexpression could not rescue the G_2_/M-phase arrest caused by Pla-B treatment. These results suggest that the decrease of cyclin B1 protein is not the cause of the G_2_/M-phase arrest caused by splicing inhibition. Since Cdk1 was phosphorylated at the Tyr15 residue in the 10 ng/ml Pla-B-treated cells (Fig. 1D), which negatively regulates Cdk1 kinase activity (28, 38, 39), overexpression of *CCNB1* might not be able to rescue the G_2_/M-phase arrest.

We investigated the effect of p27* overexpression on cell cycle progression at G_2_/M phase in detail. To this end, we established a stable cell line expressing Flag-p27* with expression controlled by a tetracycline-responsive promoter because the extensive stress of the double thymidine block and plasmid transfection affects cell cycle progression, as described above. We confirmed p27* expression only in doxycycline (DOX)-treated cells (Fig. 3G). Next, we investigated the cell cycle progression of the cells. Without DOX treatment, ∼80% of the cells exited from G_1_ phase at 8 h; conversely, ∼80% of the cells were in G_1_ phase at 14 h, suggesting that the cell cycle of the stable cell line progresses without any defect (Fig. 3H, left panel). We observed partial G_2_/M-phase arrest after DOX treatment, suggesting that p27* overexpression causes G_2_/M-phase arrest (Fig. 3H, right panel). We also investigated the expression levels of cyclin A2 in DOX-treated cells. Cyclin A2 was still observed from 14 to 20 h in DOX-treated cells (Fig. 3G), suggesting that a proportion of the Flag-p27*-expressing cells were arrested in G_2_ phase because cyclin A2 is degraded at the end of G_2_ phase (30, 32). Furthermore, we observed the morphology of the cells at 20 h and found that only ∼5% of the cells were round (Fig. 3I). This also indicates that most of the arrested cells were arrested in G_2_ phase, not in M phase.

### Knockdown of p27* partially suppresses G_2_-phase arrest caused by splicing inhibition

If the accumulation of p27* is the only cause of the G_2_-phase arrest induced by splicing inhibition, knockdown of p27* should rescue the G_2_-phase arrest. To test this hypothesis, we transfected an siRNA against *CDKN1B (p27)* into synchronized cells and harvested at 8 h after release from a double thymidine block (i.e. in G_2_/M phase). We observed only p27*, but not p27 FL, in control siRNA-treated cells after Pla-B treatment, which is consistent with the results above (Fig. 1D). In *CDKN1B* siRNA-treated cells, p27* was not observed suggesting that p27* was successfully knocked down (Fig. 4A). At 8 h, most cells were in G_2_/M phase regardless of treatment with the siRNA and Pla-B (Fig. 4B). Without Pla-B treatment, most cells transitioned to G_1_ phase at 12 h with or without siRNA treatment. Upon Pla-B treatment, the majority of cells were arrested in G_2_/M phase, which is consistent with the above results (Fig. 1B). However, even after p27* knockdown, the cells were still arrested in G_2_/M phase at 12 h. In addition, we observed the cell morphology and found that only ∼10% of the cells were round at 12 h (Fig. 4C), suggesting that these cells were arrested in G_2_ phase, but not in M phase. Therefore, p27* knockdown could not rescue the G_2_-phase arrest caused by splicing inhibition. This can be interpreted as follows: both p27* accumulation and downregulation of cell cycle regulator genes contribute to the G_2_-phase arrest caused by splicing inhibition and p27* knockdown is insufficient to rescue the G_2_-phase arrest (see discussion).

**Figure 4.**
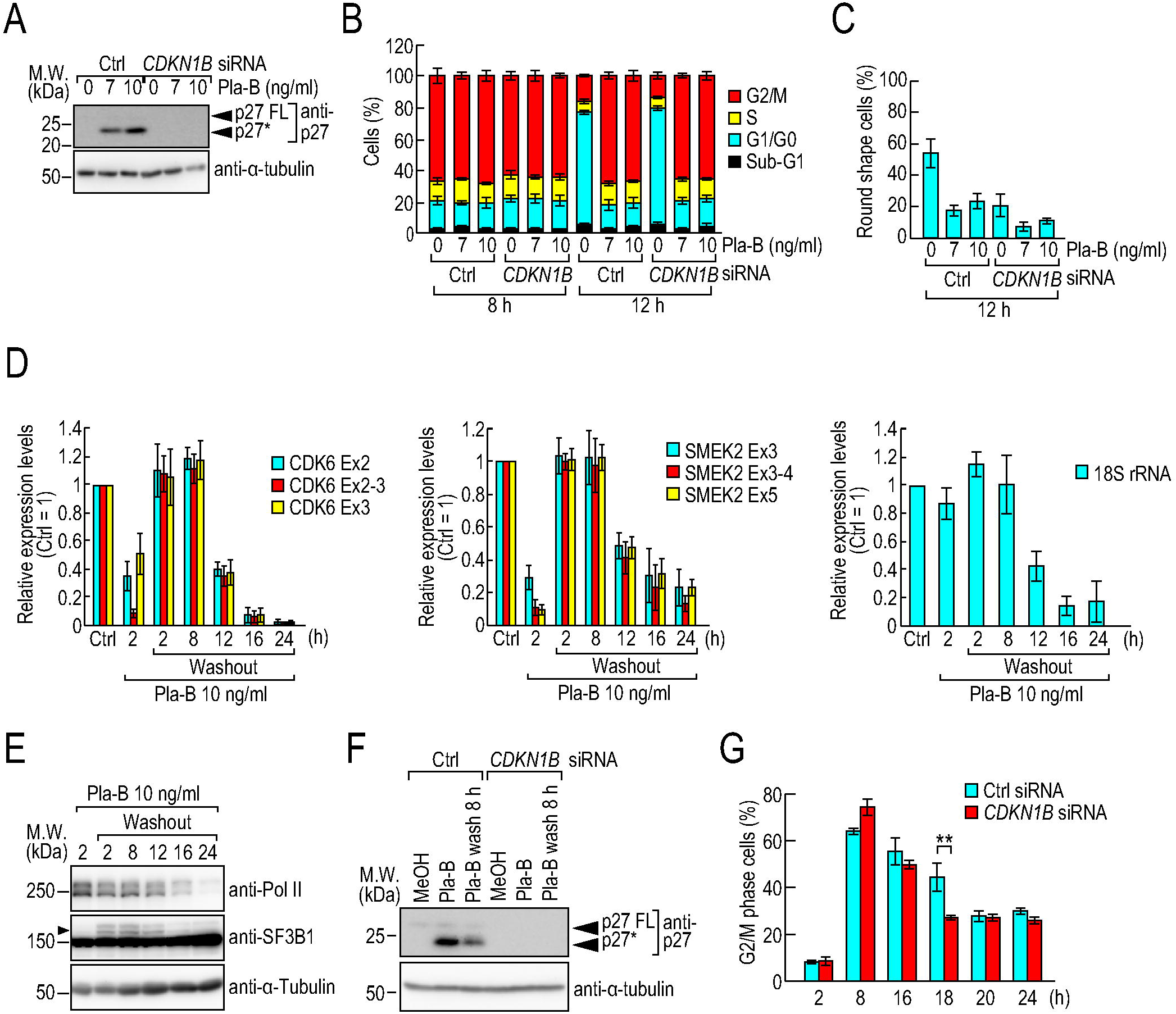
Knockdown of p27* accelerates exit from G_2_/M phase. (A) Eighteen hours after treatment with thymidine, synchronized HeLa S3 cells were transfected with control or *CDKN1B* siRNA. Two hours after release from the second thymidine block, the cells were treated with MeOH or the indicated concentration of Pla-B and cultured for an additional 6 h (G_2_/M phase), and then analyzed by immunoblotting. (B, C) HeLa S3 cells were treated with thymidine, siRNA, and MeOH or Pla-B as described in (A). Cell cycle was analyzed by cytometry at 8 or 12 h after release from the double thymidine block (B). The morphology of the cells was observed and round cells were counted at 12 h after release from the double thymidine block (C). (D, E) HeLa S3 cells were treated with 10 ng/ml Pla-B for the indicated periods and then washed out. Forty-eight hours after the addition of Pla-B, newly transcribed RNAs were labeled for 1 h and then analyzed by RT-qPCR (D). Forty-eight hours after the addition of Pla-B, protein samples were prepared and analyzed by immunoblotting using the indicated antibodies. The arrowheads indicate phosphorylated SF3B1 (E). (F) HeLa S3 cells were synchronized by a double thymidine block and transfected with *CDKN1B* siRNA. The cells were treated with 10 ng/ml Pla-B for 8 hours and then washed out. Protein levels of the indicated proteins were analyzed by immunoblotting at 24 h after the addition of Pla-B. (G) HeLa S3 cells were synchronized by a double thymidine block, treated with 10 ng/ml Pla-B for 6 h, and then washed out. The number of cells in G_2_/M phase was analyzed at the indicated time points by cytometry. Statistical significance was assessed by the two-tailed t-test (\*\**P* < 0.01). Error bars indicate s.d. (n = 3).

To segregate the effects of p27* accumulation and downregulation of cell cycle regulators, we tested whether splicing activity is recovered and p27* remains in cells after Pla-B washout. Because Pla-B binds to its target protein, SF3B1, non-covalently (21), splicing activity is supposed to be reactivated after the removal of Pla-B. After treatment of Pla-B for 2 h, the expression levels of exon-exon junctions are substantially lower than those of their adjacent exons, suggesting that Pla-B treatment inhibits splicing (Fig. 4D). After washout following 2-8 h of Pla-B treatment, splicing activity and the RNA levels of *CDK6, SMEK2* and 18S rRNA were recovered (Fig. 4D). Conversely, after treatment for 12, 16, and 24 h, the RNA levels were not recovered despite the Pla-B washout (Fig. 4D). In addition, we investigated whether SF3B1 is phosphorylated after Pla-B washout because SF3B1 is phosphorylated only in active spliceosomes (49, 50). We found that SF3B1 was not phosphorylated after Pla-B treatment for 2 h and re-phosphorylated after washout following 2-8 h of Pla-B treatment (Fig. 4E). These results suggest that splicing activity and the RNA levels were recovered after the washout following 8 h of Pla-B treatment at the longest. Next, we investigated whether p27* still remains after washout following 8 h of Pla-B treatment. After release from a double thymidine block, the cells were treated with Pla-B for 8 h and washed out, then p27* protein levels were analyzed by immunoblotting. In consistent with the above results, we observed p27*, but not p27 FL, in the Pla-B treated cells, and p27* still remained even after Pla-B washout (Fig. 4F). Therefore, we assessed the effect of p27* on cell cycle progression using the Pla-B washout cells in which splicing activity and the RNA levels were recovered and p27* still remained. We transfected synchronized cells with the control or *CDKN1B* siRNA, and treated them with Pla-B then washed out. We confirmed p27* knockdown by immunoblotting (Fig. 4F). Then, we analyzed cell cycle progression of the cells and found that the *CDKN1B* siRNA-treated cells exited from G_2_/M phase slightly but significantly earlier than control siRNA-treated cells (Fig. 4G, 18 h), indicating that p27* contributes to the G_2_/M arrest caused by splicing inhibition and its knockdown has a positive impact on cell cycle progression.

### p27* binds to and inhibits M-phase cyclins

p27* contributed to G_2_-phase arrest (Fig. 4), and therefore we investigated whether p27* inhibits the kinase activity of M-phase cyclins: Cdk1–cyclin A and Cdk1–cyclin B complexes. To investigate whether p27* bound to M-phase cyclins, we performed immunoprecipitation experiments. Flag-p27* or Flag-p27 FL was precipitated using anti-Flag tag magnetic beads and was eluted using an excess amount of Flag peptide. As a control, we tested the binding between p27 FL/p27* and components of G_1_-phase cyclin (Cdk2 and Cyclin E) because the well-known function of p27 is binding to and inhibiting G_1_-phase cyclin (41, 42). As we expected, the components of G_1_-phase cyclin were coimmunoprecipitated with Flag-p27* and Flag-p27 FL (Fig. 5A). We also found that the components of M-phase cyclins were coimmunoprecipitated with Flag-p27* and Flag-p27 FL, suggesting that p27* and p27 FL are able to bind to M-phase cyclins. Importantly, Pla-B treatment did not affect the efficiency of binding between p27 FL/p27* and cyclins (Fig. 5A). Interestingly, p27* appeared to bind to the components of G_1_- and M-phase cyclins more efficiently than p27 FL, for an unknown reason. In future study, we will investigate whether p27* binds to the cyclins more strongly than p27 FL.

**Figure 5.**
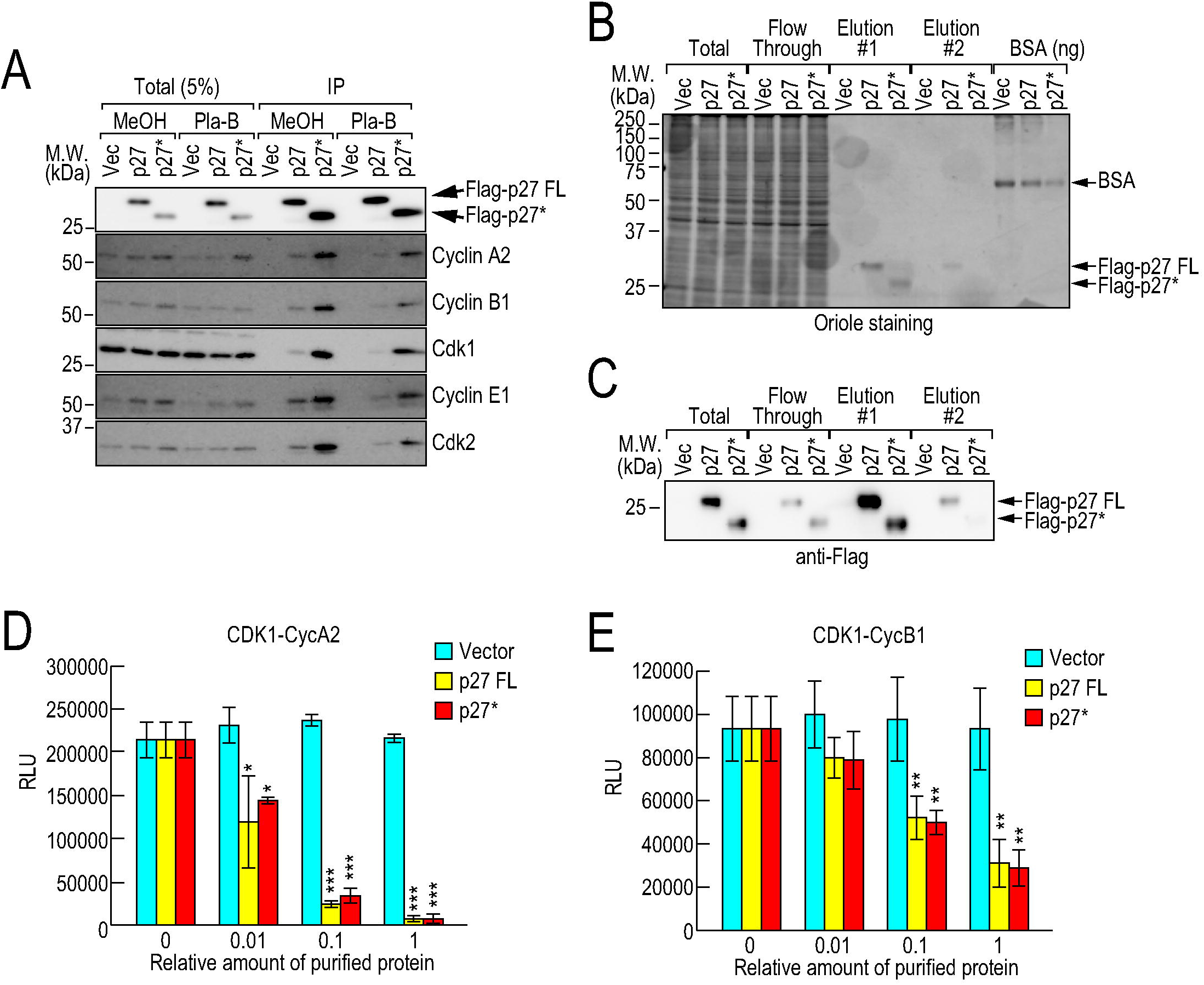
p27* binds to and inhibits M-phase cyclin. (A) HeLa S3 cells were transfected with vector, Flag-p27 FL, or Flag-p27* plasmid and treated with MeOH or 10 ng/ml Pla-B for 4 h. Immunoprecipitation was performed using anti-DDDDK (Flag) tag beads. Flag-tagged and coimmunoprecipitated proteins were eluted from the beads using a DDDDK peptide and analyzed by immunoblotting. (B, C) HEK293T cells were transfected with vector, Flag-p27 FL, or Flag-p27* plasmid, and then protein extracts were prepared. Flag-tagged proteins were purified from the protein extracts using anti-DDDDK tag beads and then eluted using a DDDDK peptide. The Flag-tagged proteins were analyzed by oriole staining (B) and immunoblotting (C). Bovine serum albumin (BSA) was also analyzed to estimate the amounts of Flag-tagged proteins. (D, E) Purified Flag-tagged proteins were applied to an *in vitro* kinase assay reaction and kinase activities of cyclin A2/Cdk1 (D) and cyclin B1/Cdk1 (E) were measured. Statistical significance was assessed by one-way ANOVA and Dunnett’s test (*: *P* < 0.05; **: *P* < 0.01; ***: *P* < 0.001). Error bars indicate s.d. (n = 3).

To investigate whether p27* inhibits the kinase activity of the CDK–cyclin complexes, we performed an *in vitro* kinase assay. Flag-tagged p27 FL and Flag-p27* were purified and successful purification of the proteins was confirmed by oriole staining and immunoblotting (Fig. 5B and 5C). Both p27 FL and p27* inhibited the kinase activity of recombinant Cdk1–cyclin A2 and Cdk1–cyclin B1 in a dose-dependent manner (Fig. 5D and 5E). These results suggest that both p27 FL and p27* bind to and inhibit M-phase cyclins.

### p27* is resistant to proteasomal degradation

We observed that p27 FL and p27* bind to and inhibit M-phase cyclins; however, only p27* was expressed in cells with G_2_ arrest caused by Pla-B treatment (Figs. 1 and 5). To investigate why only p27*, but not p27 FL, was observed in G_2_-arrested cells, we examined the splicing pattern of the *CDKN1B* gene. If splicing of *CDKN1B* pre-mRNA is completely inhibited and spliced mRNA is absent, p27 FL protein, which is translated from spliced mRNA, would not be produced. We tested this hypothesis and found that both spliced and unspliced forms of *CDKN1B* mRNA were present in the cells treated with 10 ng/ml Pla-B for 8 h, although unspliced and partially spliced forms were at much lower levels than the spliced form, presumably because these forms are degraded by the nuclear exosome and NMD (Fig. 6A). In addition, we previously found that splicing inhibition causes *CDKN1B* mRNA stabilization by unknown mechanism (23), therefore the spliced form might not have decreased even after Pla-B treatment. In either case, this result suggests that both p27 FL and p27* proteins were produced in the 10 ng/ml Pla-B-treated cells (Fig. 6A). Meanwhile, in the 3 ng/ml Pla-B-treated cells, only weak bands corresponding to unspliced or partially spliced forms were observed; therefore, p27* might not be produced in the 3 ng/ml Pla-B-treated cells, which is consistent with the results above (Fig. 1D). Interestingly, we observed no additional accumulation of pre-mRNA at 24 h. Such pre-mRNA might be degraded by the nuclear exosome and NMD. Another possibility is that transcription might be inactive at 24 h because of downregulation of transcription factors; therefore, pre-mRNA might not be produced. To test this hypothesis, we purified nascent RNA transcribed within a 1 h time window just before cell harvest (i.e., 7–8 h and 23–24 h) and analyzed it to minimize the effect of RNA degradation. We found that the spliced form of *CDKN1B* mRNA decreased while conversely the unspliced form increased upon Pla-B treatment in a dose-dependent manner at 7–8 h (Fig. 6B), suggesting that splicing was inhibited by Pla-B. Interestingly, both spliced and unspliced forms decreased at 23–24 h (Fig. 6B), suggesting that transcription is inactive, so consequently no accumulation of the unspliced form.

**Figure 6.**
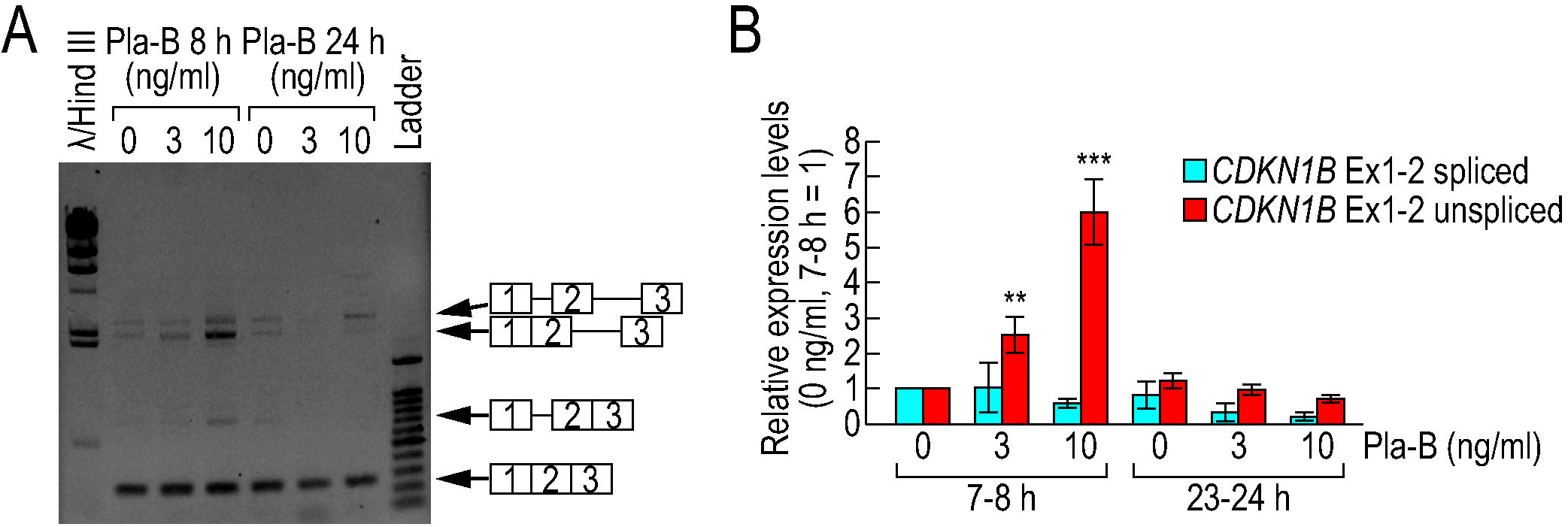
Both spliced and unspliced forms of *CDKN1B* mRNA were observed in Pla-B-treated cells. (A) Two hours after release from a double thymidine block, synchronized HeLa S3 cells were treated with MeOH or the indicated concentrations of Pla-B. The cells were harvested at 8 or 24 h after release from this block and total RNA was purified. Splicing pattern changes were analyzed by RT-PCR. (B) Two hours after release from a double thymidine block, synchronized HeLa S3 cells were treated with MeOH or the indicated concentrations of Pla-B. At 7 or 23 h after release from the block, the cells were treated with 200 μM ethynyl uridine and cultured for 1 h to label the newly synthesized RNA. The newly synthesized RNA was purified and the relative amounts of spliced and unspliced forms of *CDKN1B* mRNA were analyzed by qRT-PCR. Statistical significance was assessed by one-way ANOVA and Dunnett’s test (*: *P* < 0.05; **: *P* < 0.01; ***: *P* < 0.001). Error bars indicate s.d. (n = 3).

The above results suggest that both p27 FL and p27* are produced in the G_2_-arrested cells. Therefore, we assumed that p27 FL is degraded by the proteasome in G_2_-arrested cells, but p27* is not. Indeed, in the C-terminus of p27 FL, there is a phosphorylation site (threonine 187), the phosphorylation of which is the trigger for ubiquitylation and proteasomal degradation (51). This mechanism keeps the p27 FL protein level low in G_2_/M phase. In contrast, p27* lacks this phosphorylation site because of the C-terminus truncation and is not supposed to be ubiquitinated and degraded by the proteasome in G_2_/M phase (15). However, there is no experimental evidence of the hypothesis. To investigate whether p27* escapes proteasomal degradation, we evaluated the expression levels of Flag-p27 FL and Flag-p27* in G_2_/M phase by synchronizing the cell cycle of Flag-p27 FL- and Flag-p27*-expressing cells with expression controlled by DOX. Even after DOX treatment, Flag-p27 FL was barely expressed and the amount of Flag-p27 FL was increased by MG132, a potent proteasome inhibitor, suggesting that Flag-p27 FL was degraded by the proteasome in G_2_/M phase (Fig. 7A), which is consistent with a previous report (44). However, the amount of Flag-p27* was not increased by MG132 treatment. We also investigated the proteasomal degradation of endogenous p27 FL and p27*. The cell cycle of HeLa cells was synchronized, and the cells were treated with Pla-B and MG132. We found that the amount of endogenous p27 FL was increased by MG132 in G_2_/M phase, but that of p27* was not, suggesting that endogenous p27* is also resistant to proteasomal degradation in G_2_/M phase (Fig. 7B). These results strongly suggest that p27 FL is degraded by the proteasome in G_2_/M phase, but p27* is not. Because p27* does not have the phosphorylation site for ubiquitylation, we compared the ubiquitylation levels of p27 FL and p27*. To investigate the ubiquitylation level, HEK293T cells were transfected with Flag-p27 FL, Flag-p27*, or vector plasmids. To observe ubiquitylation of p27 FL and p27* clearly, the cells were treated with MG132 and transfected with the HA-Skp2 plasmid (52) because Skp2 protein recruits p27 to SCF ubiquitin ligase and contributes to the degradation of p27 in G_2_/M phase. Ubiquitylated proteins were purified from the cells using Tandem Ubiquitin-Binding Entity (TUBE) (53) and analyzed by immunoblotting (Fig. 7C). We observed stronger signals of higher-molecular-weight bands of Flag-p27 FL than those of Flag-p27*. This indicates that p27* was less ubiquitylated by the SCF ubiquitin ligase than p27 FL, presumably because p27* lacks the phosphorylation site to trigger ubiquitylation by the SCF complex. As a control, we also investigated the ubiquitylation level of p27 FL and p27* by the Kip1 ubiquitylation promoting complex (KPC), which ubiquitylates p27 in G_1_ phase (54, 55). This ubiquitylation is independent of T187 phosphorylation (56). In KPC-overexpressing cells, p27* was ubiquitylated to the same extent as p27 FL, suggesting that both proteins are ubiquitylated and degraded by the proteasome in G_1_ phase (Fig. 7D). Taken together, these results suggest that p27 FL is degraded by the ubiquitin–proteasome pathway in G_2_/M phase, but p27* is resistant to degradation; consequently, only p27*, but not p27 FL, accumulates at G_2_/M phase and contributes to G_2_/M-phase arrest in Pla-B-treated cells.

**Figure 7.**
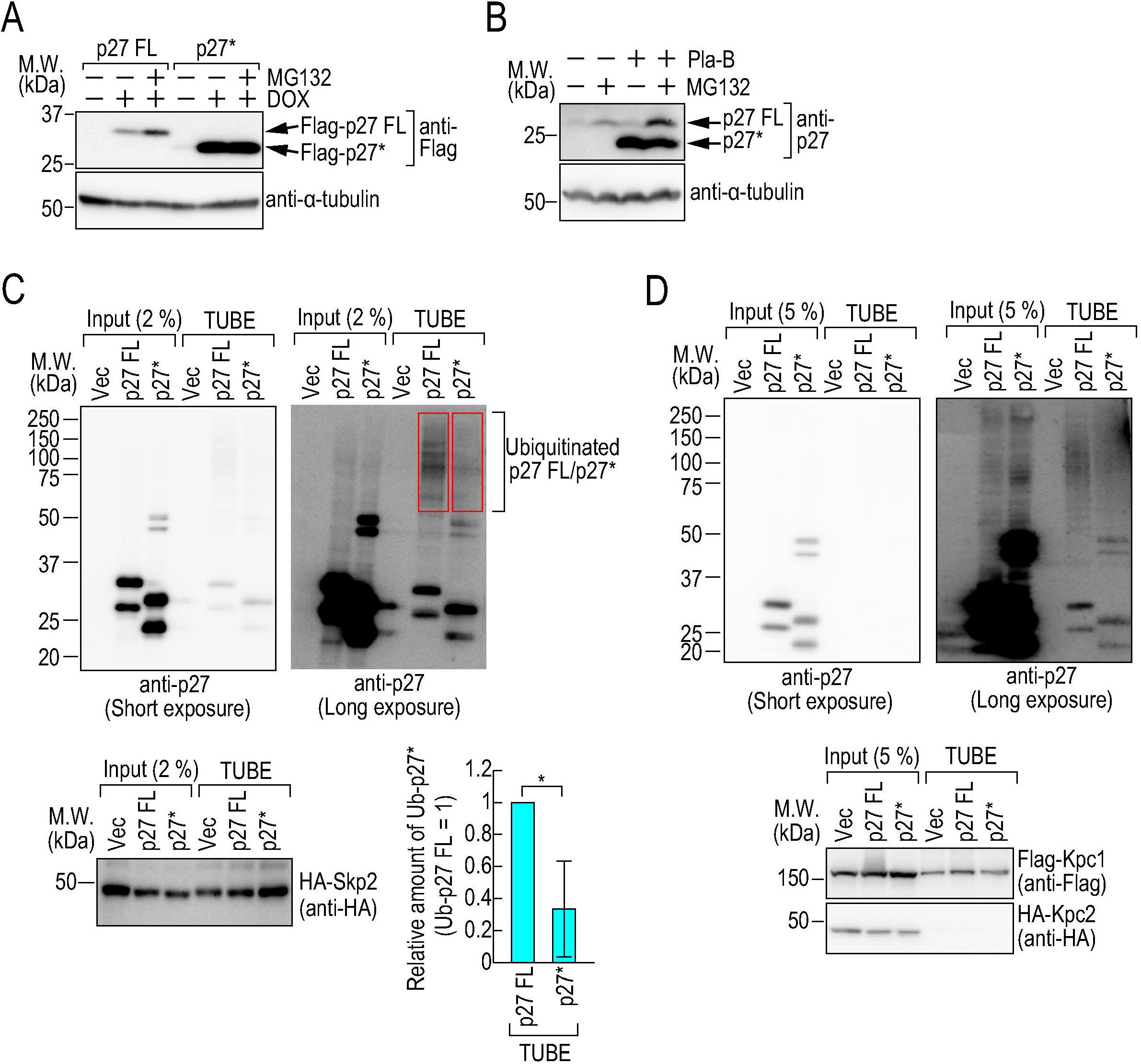
p27* is resistant to proteasomal degradation. (A) Cells expressing Flag-p27 FL or Flag-p27* under the control of tetracycline were synchronized by a double thymidine block. The cells were treated with 2 μg/ml doxycycline (DOX) at the same time as the second thymidine treatment. After release from the double thymidine block, the cells were treated with 2 μg/ml DOX and 10 μM MG132 for 8 h (G_2_/M phase) and levels of the indicated proteins were analyzed by immunoblotting. (B) Two hours after release from a double thymidine block, synchronized HeLa S3 cells were treated with MeOH or 10 ng/ml Pla-B. Six hours after release from this block, the cells were treated with 20 μM MG132. Eight hours after release from this block (G_2_/M phase), protein levels of indicated proteins were analyzed by immunoblotting. (C) HEK293T cells were transfected with HA-Skp2 and Flag-p27 FL or Flag-p27* and then treated with 20 μM MG132 for 4 h. Cell extracts were incubated with Agarose-TUBE for 4 h to purify ubiquitylated proteins. Purified proteins were analyzed by immunoblotting. The relative band intensities (p27 FL = 1) were quantified and are shown in the bar graph (bottom right panel). Red rectangles indicate the areas where band intensity was measured. Statistical significance was assessed by the two-tailed t-test (\**P* < 0.05). Error bars indicate s.d. (n = 3). (D) HEK293T cells were transfected with Flag-KPC1, HA-KPC2, and Flag-p27 FL or Flag-p27*, and then treated with 10 μM MG132 for 4 h. Cell extracts were incubated with Agarose-TUBE for 4 h to purify ubiquitylated proteins. Purified proteins were analyzed by immunoblotting.

## Discussion

Pre-mRNA splicing is one of the most important mechanisms for transcriptome integrity and proteome integrity in eukaryotes. Abnormalities in this process might cause the production of aberrant, non-functional proteins translated from pre-mRNA, which might inhibit functional proteins. To prevent the production of such abnormal proteins, cells have several mRNA quality control mechanisms (10-14): pre-mRNA degradation by the nuclear exosome, nuclear retention of pre-mRNA, and degradation of unspliced and partially spliced mRNAs by NMD. We reported that very few selected pre-mRNAs with introns increase in the cytoplasm under splicing-deficient conditions (57). This indicates that the mRNA quality control mechanisms indeed prevent the accumulation of most pre-mRNAs in the cytoplasm under splicing-deficient conditions. Exceptionally, we found that *CDKN1B (p27)* pre-mRNA increases in the cytoplasm under splicing-deficient conditions (57). However, as described above, most of the unspliced and partially spliced forms of *CDKN1B* mRNA appeared to be degraded by the nuclear exosome and NMD. Small population of unspliced and partially spliced forms might escape from the nuclear retention mechanism, because *CDKN1B* pre-mRNA (5.2 kb) is much shorter than the average length of human genes (27 kb) (58). We have revealed that short pre-mRNAs with a weak 5’
s splice site tend to escape from the nuclear retention mechanism (57). Therefore, *CDKN1B* pre-mRNA might be able to escape from the retention mechanism.

Once p27* is produced from a small quantity of remaining *CDKN1B* pre-mRNA, p27* appears to remain in splicing-deficient cells for a long time. In this study, we observed that p27* is resistant to proteasomal degradation in G_2_/M phase, presumably because it does not possess the site the phosphorylation of which is the trigger for ubiquitination from S to M phase (44, 51). Therefore, p27* is highly stable compared with p27 FL. In contrast, p27 FL was barely observed in G_2_/M phase without MG132 (Fig. 7). Another feature might also make p27* a stable protein. p27 FL consists of 198 amino acids, among which exon 1 of *CDKN1B* encodes 158 (15). Most of p27* is thus identical to p27 FL. Therefore, p27* does not appear to be a short peptide, but rather a stable protein. In addition to stability, the high identity between p27* and p27 FL indicates that p27* has physiological function. The CDK inhibitory domain of p27 resides in the N-terminus of p27 (41), which is also included in p27*. As a consequence, p27* has the same potent CDK inhibitory activity as p27 FL. Taken together, these features of *CDKN1B* pre-mRNA and p27* protein make p27* a stable protein with a physiological function.

As described above, the overexpression of p27* caused G_2_-phase arrest. However, knockdown of p27* could not rescue the G_2_-phase arrest caused by splicing inhibition. We assume that both p27* accumulation and downregulation of cell cycle regulators contribute to the G_2_-phase arrest caused by splicing inhibition. Indeed, it was reported that several cell cycle regulator genes including *CCNA2, PLK1, PLK2, WEE1, AURKA*, and *AURKB* showed splicing pattern changes and decreases in mRNA level in splicing-inhibited cells (19, 57). Therefore, p27* knockdown is insufficient to rescue the G_2_-phase arrest, and both p27* knockdown and upregulation of the cell cycle regulators might be required to rescue the G_2_-phase arrest caused by splicing inhibition. In addition, the decrease of cell cycle regulators might explain the result of the washout experiment (Fig. 4G). After the washout of Pla-B, gene expression would recover and such cell cycle regulators might start to accumulate for cell cycle progression. When cell cycle regulators accumulate sufficiently to overcome the inhibitory activity of p27* in the cells, the cells could transit from G_2_/M phase to G_1_ phase. If p27* is knocked down, the threshold of the amount of cell cycle regulator required for cell cycle progression must become lower because of a lack of inhibitory activity. Therefore, cells with p27 knockdown might exit G_2_/M phase slightly but significantly earlier than control cells (Fig. 4G).

Why do cells produce p27* under splicing-deficient conditions? We speculate that the *CDKN1B* gene functions as a sensor for splicing abnormalities. The transcriptome and proteome are perturbed in splicing-deficient cells, and such perturbation might cause splicing-related diseases, including myelodysplastic syndromes and leukemia (5-9). Therefore, the proliferation of splicing-deficient cells might increase the risk of such splicing-related diseases. To reduce this risk, the proliferation of splicing-deficient cells should be inhibited. To inhibit the proliferation of splicing-deficient cells, p27* is a suitable protein because this stable protein with cell cycle inhibitory activity is translated from pre-mRNA only under splicing-deficient conditions. We assume that this system protects the body from splicing abnormality by minimizing the effects of splicing-deficient cells. We also speculate that there are other truncated proteins translated from pre-mRNA to adapt to splicing-deficient conditions. Indeed, we found that another truncated protein contributed to cell death upon splicing inhibition (unpublished data). In a future study, we plan to reveal the entire molecular mechanism by which cells adapt to splicing-deficient conditions using truncated proteins.

The system by which truncated proteins inhibit cell proliferation can be adopted for efforts to achieve tumor suppression. If we inhibit the splicing of cancer cells, the proliferation of such cells should be inhibited by the truncated proteins and downregulation of gene expression. Indeed, Pla-B and SSA have been reported to be potent antitumor reagents (15-18, 59, 60). Although a phase I clinical trial of E7017—a derivative of Pla-B—was discontinued because of serious side effects (61), H3B-8800—another Pla-B derivative—has been investigated in a phase 1/2 study (60, 62). Thus far, the compound has been developed for use on myelodysplasia, acute myelogenous leukemia, and chronic myelomonocytic leukemia patients with spliceosome mutations. This study and future studies regarding truncated proteins translated from pre-mRNA might contribute to the development of a novel antitumor reagent with fewer side effects that targets a broad range of cancers.

In this study, we found that p27* is produced from pre-mRNA in splicing-deficient cells and inhibits cell cycle progression through binding to and inhibiting M-phase cyclins. We believe that these findings deepen our understanding of the mechanisms that protect the body from splicing abnormality and could aid the development of novel anticancer drugs based on splicing inhibitors.

## Materials and Methods

### Cell culture and synchronization

HeLa S3 and HEK293T cells were cultured in Dulbecco’s modified Eagle’s medium containing 10% heat-inactivated fetal bovine serum (Thermo Fisher Scientific, Waltham, MA, USA). Cells were maintained in 5% CO_2_ at 37°C. For cell cycle synchronization, the cells were treated with 2 mM thymidine (FUJIFILM Wako Pure Chemical Corporation, Osaka, Japan) for 18 h. After treatment, the cells were washed twice with phosphate-buffered saline (PBS) to release them from the thymidine block and then cultured in fresh culture medium for 8 h. The cells were treated with 2 mM thymidine again for 16 h and then washed with PBS twice to release them from the double thymidine block.

### Antibodies and reagents

A mouse monoclonal anti-α-tubulin antibody (T6074) was purchased from Sigma-Aldrich (St. Louis, MO, USA). Mouse monoclonal anti-cyclin A2 (#4656), rabbit polyclonal anti-cyclin B1 (#4138), mouse monoclonal anti-cdc2 (Cdk1) (#9116), rabbit polyclonal anti-phospho-cdc2 (Tyr15) (#9111), mouse monoclonal anti-cyclin E1 (#4129), and rabbit monoclonal anti-p27 (#3686) antibodies were purchased from Cell Signaling Technology (Danvers, MA, USA). Mouse monoclonal anti-RNA polymerase II antibody (8WG16) was purchased from BioLegend (San Diego, CA, USA). Mouse monoclonal anti-Sap155 (SF3B1) (D221-3), mouse monoclonal anti-DDDDK (Flag) (M185-3L), rabbit polyclonal anti-HA (561), mouse monoclonal anti-Myc (M192-3) antibodies, and anti-DDDDK beads (M185-11R) were purchased from Medical and Biological Laboratories (MBL) (Nagoya, Japan). Rabbit polyclonal anti-PARP-1 (H-250) and mouse monoclonal anti-Cdk2 (D-12) antibodies were purchased from Santa Cruz Biotechnology (Santa Cruz, CA, USA). HRP-conjugated anti-mouse IgG and anti-rabbit IgG secondary antibodies were purchased from GE Healthcare (Chicago, IL, USA). Doxycycline, puromycin, and G418 were purchased from TaKaRa Bio (Otsu, Japan). Pladienolide B was purchased from Santa Cruz Biotechnology. MG132 was purchased from Sigma-Aldrich.

### Cell count and cell viability assay

After trypsinization, the numbers of viable and dead cells were counted using trypan blue exclusion and a hemocytometer.

### Cell cycle analysis

Cells were fixed in 70% ethanol, rinsed with PBS, and then stained with a solution containing 20 μg/ml propidium iodide (Thermo Fisher Scientific), 0.05% Triton X-100, and 0.1 mg/ml RNase A (Thermo Fisher Scientific). The cell cycle was monitored by the image-based cytometer Tali (Thermo Fisher Scientific).

### Morphological observation

Cells were synchronized using thymidine and treated with Pla-B. Morphology of the cells was then observed under an IX73 microscope (Olympus, Tokyo, Japan).

### RNA preparation and RT-PCR

Total RNA was extracted from cells using TRIzol reagent (Thermo Fisher Scientific), following the manufacturer’s instructions. cDNA was prepared using Primescript Reverse Transcriptase (TaKaRa Bio) and random primers. PCR was performed using TaKaRa ExTaq (TaKaRa). PCR products were analyzed using a 1% agarose gel. To measure splicing activity, we purified nascent RNA using the Click-iT Nascent RNA Capture Kit (Thermo Fisher Scientific). Briefly, cells were treated with 200 μM 5-ethynyl-uridine for 1 h and total RNA was extracted from cultured cells using TRIzol reagent. Labeled RNA was biotinylated by the click reaction and biotinylated RNA was purified using streptavidin beads. For quantitative RT-PCR, cDNA was synthesized using the SuperScript VILO cDNA Synthesis Kit (Thermo Fisher Scientific). Quantitative RT-PCR and relative quantification analyses were performed with the MX3000P system (Agilent, Santa Clara, CA, USA) using SYBR Green dye chemistry. All primers are listed in Table S1.

### Immunoblotting

Cells were directly lysed on plates with 1× SDS-PAGE sample buffer. Proteins were then separated by SDS-PAGE. After electrophoresis, the proteins were transferred onto a PVDF membrane by electroblotting. Following incubation of the membrane with primary and secondary antibodies using standard techniques, protein bands were detected using a NOVEX ECL Chemiluminescent Substrate Reagent Kit (Thermo Fisher Scientific) on an ImageQuant LAS 4000mini (GE Healthcare).

### Plasmids and plasmid transfection

p27-int-HA plasmid was as described previously (15). Flag-p27 and Flag-p27* plasmids were as described previously (22). Flag-KPC1 and HA-KPC2 were kind gifts from Dr. Keiichi Nakayama (Kyushu University) (54). HA-Skp2 plasmid was a kind gift from Dr. Yukiko Yoshida (Tokyo Metropolitan Institute of Medical Science) (52). To construct Flag-*CCNB1* plasmid, the DNA fragment of *CCNB1* was amplified by PCR from a cDNA library of HeLa S3 cells using hCCNB1 cloning F and hCCNB1 cloning R primers. The PCR product was digested with *Bam* HI and *Not* I and subcloned into pcDNA3.1-FLAG (22). The primers used for plasmid construction are listed in Table S1. Plasmid transfection was performed using Lipofectamine 3000 Reagent (Thermo Fisher Scientific), in accordance with the manufacturer’s instructions.

### Stable cell line establishment

To establish HeLa cells expressing Flag-p27 FL or Flag-p27* under the control of tetracycline, a DNA fragment of GFP was amplified by PCR from pcDNA6.2 emGFP using GFP Hind III-Eco RI F and GFP ATTTA R primers. The PCR product was digested with EcoRI and KpnI, and subcloned into pTRE3G-BI (TaKaRa Bio) to construct pTRE3G-BI-GFP. The DNA fragments of Flag-p27 FL and Flag-p27* were amplified from Flag-p27 and Flag-p27* plasmids (22), respectively. To amplify Flag-p27, FLAG Hind III-Bgl II F and hp27 cloning R primers were used. To amplify Flag-p27*, FLAG Hind III-Bgl II F and hp27* cloning R primers were used. The PCR products were digested with *Bgl* II and *Not* I, and subcloned into pTRE3G-BI-GFP to construct pTRE3G-BI-GFP-FLAG-p27 FL and pTRE3G-BI-GFP-FLAG-p27*. HeLa Tet On 3G cells (TaKaRa Bio) were transfected with pTRE3G-BI-GFP-FLAG-p27 FL or pTRE3G-BI-GFP-FLAG-p27*, and stable clones were selected by puromycin (1 µg/ml) treatment followed by clonal selection. The primers used for plasmid construction are listed in Table S1.

Stable clones were grown in medium containing Tet system-approved fetal bovine serum (TaKaRa Bio), puromycin, and G418. To induce expression, cells were treated for the indicated times with doxycycline (2 μg/ml).

### siRNA transfection

Silencer select p27/CDKN1B siRNA (s2837 cat# 4390824) was purchased from Thermo Fisher Scientific. siGENOME Control Pool Non-Targeting #2 (cat# D-001206-14-20) was purchased from GE Healthcare. siRNA transfection was performed using Lipofectamine RNAiMAX (Thermo Fisher Scientific), in accordance with the manufacturer’s instructions.

### Immunoprecipitation

Cells were suspended in lysis buffer [25 mM HEPES, pH 7.5, 150 mM NaCl, 2 mM MgCl_2_, 1 mM EDTA, 2.5 mM EGTA, 1% Nonidet P-40, 10% glycerol, cOmplete Protease Inhibitor cocktail (Sigma-Aldrich), and PhosSTOP (Sigma-Aldrich)] and sonicated for 15 s, and then incubated on ice for 15 min. After centrifugation, cell extracts were incubated with anti-DDDDK-tag beads for 1 h at 4°C with gentle agitation. The beads were washed three times with lysis buffer, and then the bound proteins were eluted using DDDDK-tag peptides (MBL; 3325-205). The eluted proteins were mixed with 1× SDS-PAGE sample buffer and heated at 95°C for 5 min.

For the purification of ubiquitylated proteins, cells were suspended in lysis buffer [25 mM HEPES, pH 7.5, 150 mM NaCl, 2 mM MgCl_2_, 1 mM EDTA, 2.5 mM EGTA, 1% Nonidet P-40, 10% glycerol, cOmplete Protease Inhibitor cocktail (Sigma-Aldrich), PhosSTOP (Sigma-Aldrich), and 1 mM N-ethylmaleimide (Nacalai Tesque, Kyoto, Japan)], sonicated for 15 s, and then incubated on ice for 15 min. After centrifugation, cell extracts were incubated with Agarose-TUBE2 (LifeSensors, Malvern, PA, USA) at 4°C for 4 h with gentle agitation. The beads were washed three times with lysis buffer, and then the bound proteins were extracted with 1× SDS-PAGE sample buffer by heating at 95°C for 5 min.

### *In vitro* kinase assay

HEK293T cells were transfected with vector, Flag-p27, or Flag-p27* plasmids using Lipofectamine 3000 Reagent, in accordance with the manufacturer’s instructions. Lysates of the transfected cells were prepared using lysis buffer (20 mM HEPES-KCl pH 7.4, 150 mM NaCl, 2 mM EDTA, 1% Triton, and cOmplete Protease Inhibitor cocktail). Flag-tagged proteins were purified from the cell lysates using a DDDDK-tagged protein purification kit (MBL), in accordance with the manufacturer’s instructions. To evaluate the purification efficiency, the purified proteins were separated by SDS-PAGE, stained with Oriole Fluorescent Gel Stain (Bio-Rad), in accordance with the manufacturer’s instructions, and analyzed by immunoblotting. The *in vitro* kinase assay was performed using the Cyclin A2/Cdk1 Kinase Enzyme System (Promega, Madison, WI, USA), recombinant Cyclin B1/Cdk1 (SignalChem Biotech Inc., Richmond, BC, Canada), and ADP-Glo kinase assay (Promega), in accordance with the manufacturers’ instructions. Briefly, the recombinant Cdk1–cyclin A2 or Cdk1–cyclin B1 was mixed with the purified Flag-tagged proteins and then incubated at 25°C for 10 min. Histone H1 and ATP were added to the reaction, followed by incubation at 25°C for 1 h. Next, ADP-Glo Reagent was added to the reaction, followed by incubation at 25°C for 40 min. Finally, Kinase Detection Reagent was added to the reaction, followed by incubation at 25°C for 30 min. Luminescence was measured using a Varioskan Flash (Thermo Fisher Scientific).

### Statistical analysis

Statistical analysis was performed using R Commander. A two-tailed t-test (Fig. 3B, 4G, and 7C), one-way ANOVA with Dunnett’s test (Fig. 5D and 5E), and one-way ANOVA with Tukey’s test (Fig. 3D) were performed to determine statistical significance. Data are presented as means ± standard deviation. The sample size used in each experiment is stated in the figure legends. P < 0.05 was considered statistically significant.

**Table 1.**
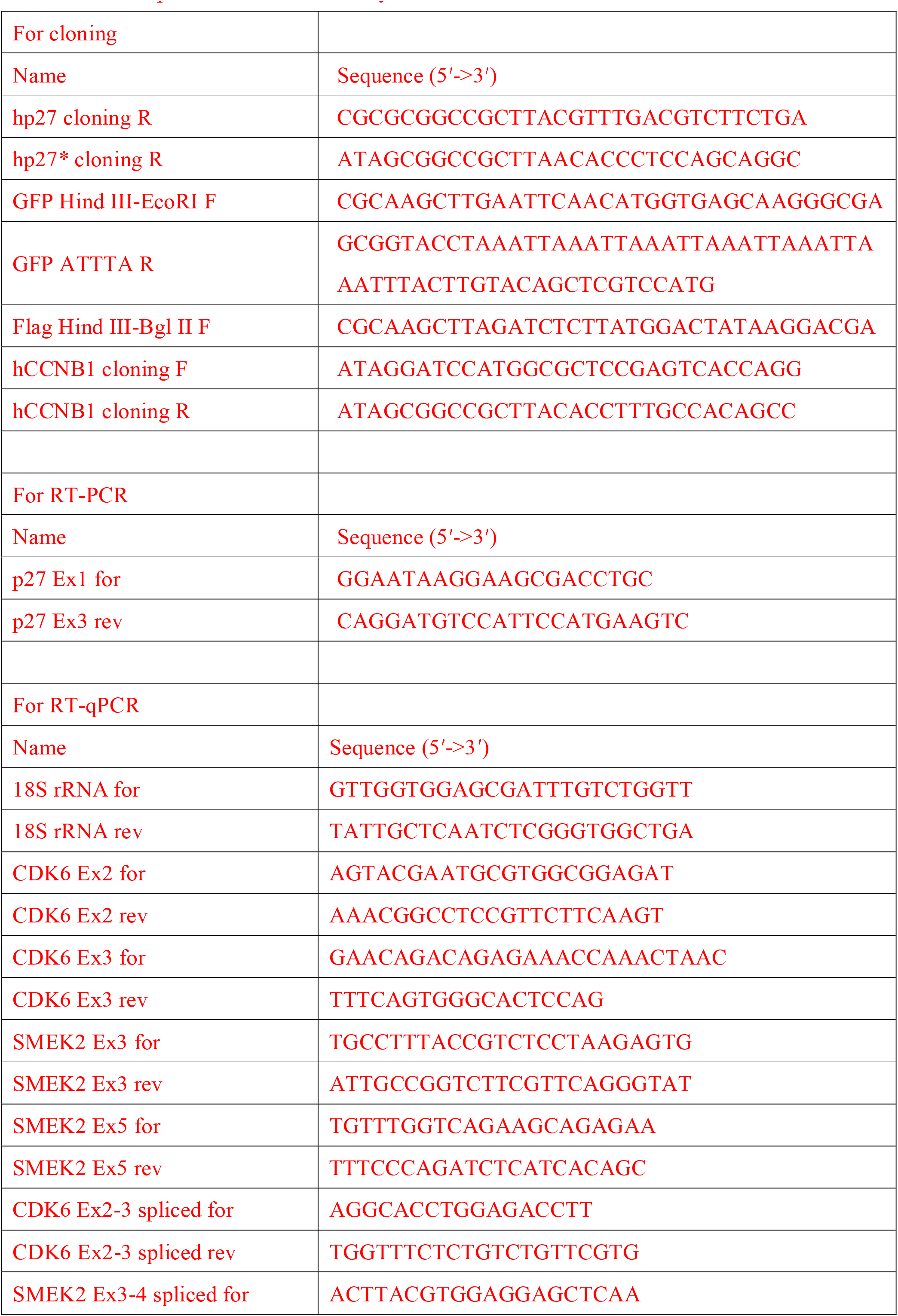
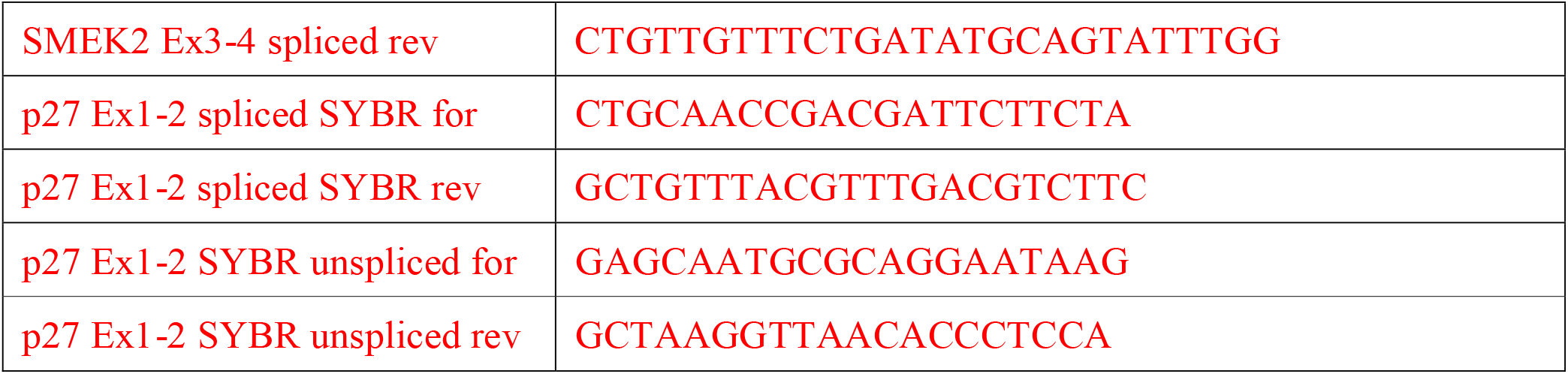
List of primers used in this study.

## Data Availability

The data underlying the results are available upon request.

## Acknowledgments

We are grateful to Mr. Kei Kikuchi, Ms. Midori Shima, Ms. Shiori Nakada, and Mr. Kanjiro Nakagawa for technical assistance. We thank Dr. Yukiko Yoshida (Tokyo Metropolitan Institute of Medical Science) for HA-Skp2 plasmid. Additionally, we thank Dr. Keiichi Nakayama (Kyushu University) for Flag-KPC1 and HA-KPC2 plasmids. This research was funded by JSPS KAKENHI (JP21H02423 to D.K.), Takeda Science Foundation (to D.K.), Tamura Science & Technology Foundation (to D.K.), the Kato Memorial Bioscience Foundation (to D.K.), The Ichiro Kanehara Foundation (to D.K.), and Suzuken Memorial Foundation (to D.K.). Finally, we thank Edanz (https://jp.edanz.com/ac) for editing a draft of this manuscript.

